# CoRhythMo: A Computational Framework for Modeling Biobehavioral Rhythms from Mobile and Wearable Data Streams

**DOI:** 10.1101/2020.08.10.244020

**Authors:** Runze Yan, Xinwen Liu, Janine M Dutcher, Michael J Tumminia, Daniella Villalba, Sheldon Cohen, David Creswell, Kasey Creswell, Jennifer Mankoff, Anind K Dey, Afsaneh Doryab

## Abstract

This paper presents CoRhythMo, the first computational framework for modeling biobehavioral rhythms - the repeating cycles of physiological, psychological, social, and environmental events - from mobile and wearable data streams. The framework incorporates four main components: mobile data processing, rhythm discovery, rhythm modeling, and machine learning. We use a dataset of smartphone and Fitbit data collected from 138 college students over a semester to evaluate the framework’s ability to 1) model biobehavioral rhythms of students, 2) measure the stability of their rhythms over the course of the semester, 3) model differences between rhythms of students with different health status, and 4) predict the mental health status in students using the model of their biobehavioral rhythms. Our evaluation provides evidence for the feasibility of using CoRhythMo for modeling and discovering human rhythms and using them to assess and predict different life and health outcomes.

## Introduction

The term biobehavioral rhythms introduced in [1], refers to the repeating cycles of physiological (*e*.*g*., heart rate and body temperature), psychological (*e*.*g*., mood), social (*e*.*g*., work events), and environmental (*e*.*g*., weather) that affect human body and life. Rooted in Chronobiology, “the scientific discipline that quantifies and explores the mechanisms of biological time structure and their relationship to the rhythmic manifestations in the living matter” [2], biobehavioral rhythms aim at studying cyclic events observed in human data collected from personal and consumer level mobile and wearable devices [1]. Such devices provide the capability of continuous tracking of biobehavioral signals of individuals in their daily life and outside of controlled lab settings which have been the standard method for studying biological rhythms.

Numerous research studies have shown the impact of understanding rhythms and their effect on human life and wellbeing. For example studies in [1, 3, 4] demonstrate the association between long-term disruption in biological rhythms and health outcomes such as cancer, diabetes, and depression. Other studies have shown the impact of shift work on quality of life in shift workers such as nurses and doctors [5, 6]. These studies, however, have often been limited to controlled settings to observe certain behaviors and effects. With passive sensing of physiological and behavioral signals from mobile and wearable devices, it is now possible to study human rhythms more broadly and holistically in the wild through the collection of biobehavioral data from different sources. This opportunity, however, introduces new challenges. First, the longitudinal time series data collected from personal devices is massive, noisy, and incomplete requiring careful processing to extract and preserve useful fine-grained knowledge from data in various temporal granularity levels to be used for further modeling. Second, the fact that each data source (*e*.*g*., smartphone sensors) can capture different aspects of human rhythms (biological, behavioral or both) requires exploration and incorporation of each signal to identify biological and behavioral indicators on the micro and macro level that may reveal a cyclic behavior. This process can be exhaustive and needs automation. Moreover, although the modeled rhythms by themselves can provide useful insights into human health and life, the exhaustive number of rhythm models generated by each source makes it difficult for manual interpretation of the models by researchers or experts. A further computational step should incorporate those models to provide further insights into different health and lifestyle outcomes both physical and mental.

We propose a computational framework to address the aforementioned challenges through a series of data processing and modeling steps. The framework first processes the raw sensor data collected from mobile and wearable devices to extract high-level features from those data streams. It then models biobehavioral rhythms for each sensor feature alone and in combination with other features to discover rhythmicity and other characteristics of cyclic behavior in the data. The biobehavioral rhythm models provide a series of characteristic features which are further used for measuring stability in biobehavioral rhythms and to predict different outcomes such as health status through a machine learning component. We evaluate the framework using mobile and Fitbit data collected from 138 college students over a semester to test the framework’s ability to detect rhythmicity in students’ data in different time frames over the course of the semester and to measure the stability and variation of rhythms among students with different mental health status. We then use the models of the rhythms to predict the mental health status of students at the end of the semester. Our research makes the following contributions:

1. We introduce the first computational framework for modeling biobehavioral rhythms that provides the ability to
  a. flexibly process massive sensor data in different time granularity thus providing the ability to model and observe short- and long-term rhythmic behavior.
  b. identify variation and stability in individual and groups of time series data.
  c. help observe the impact of cyclic biobehavioral parameters in revealing and predicting different outcomes (*e*.*g*., health).
2. We introduce two novel methods for measuring rhythm stability that incorporate the autocorrelation sequences and Cosinor models to generate stability scores in different time segments and across different populations.
3. We demonstrate the framework’s ability to explore and discover biobehavioral rhythms in college students data and highlighting differences in rhythms among students with different mental health status.

In the following sections, we describe related work in the domain of mobile health and behavior modeling and discuss the motivation for modeling cyclic human behavior and its potential role in revealing health status. We then present our computational framework followed by the case study in modeling biobehavioral rhythms in college students and explore the role of those models in predicting students’ mental health status. In particular, we look at two long-term mental health conditions, namely depression and loneliness. Our analysis and results provide detailed insights into the relationship between biobehavioral rhythms and mental health status in students. We discuss those insights and their implications for building rhythm-aware technology for health and well-being.

## Background and Related Work

### Biological rhythms

The assessment of rhythmic phenomena in living organisms reveals the existence of events and behavior that repeat themselves in certain cycles and can be modeled with periodic functions [2, 7]. Each periodic function is specified by its average level, oscillation degree, and time of oscillation optimal. Biological rhythms, including patterns of activity and rest or circadian rhythms have been extensively studied in Chronobiology and medicine [1, 3, 4] mostly in controlled environmental settings.

The advancements in activity trackers have made it possible to study these phenomena outside of the labs and have demonstrated the reliability of such devices in capturing circadian disruptions, including sleep and physical and mental health conditions. For example, studies using research-grade actigraphy devices have shown differences in circadian rhythms among patients with bipolar disorder, ADHD, and schizophrenia [8]. Other studies have used the same type of data to explore circadian disruption in cancer patients undergoing chemotherapy [8, 9]. Commercial devices such as Fitbits are now able to infer sleep duration and quality reasonably accurately. Two brief studies with healthy young adults have used activity data from Fitbit devices to quantify rest-activity rhythms and found that rhythm measurement compared well relative to research-grade actigraphy [10, 11]. Studies in [12] and [13] have also explored the capability of personal tracking devices to measure sleep compared to gold standards such as polysomnography.

### Behavior Modeling in the Wild via Mobile Sensing

The study of biobehavioral rhythms also relates to the research in understanding human behavior from passive sensing data collected via smartphones and wearable devices. Only a few studies have actually used mobile data for understanding the circadian behavior of different chronotypes (*e*.*g*., [14–16]). Abdullah et al. [14] analyzed the patterns of phone usage to demonstrate differences in the sleep behavior of early and late chronotypes. In a similar study using the same type of data, they showed the capability of using mobile data to explore daily cognition and alertness [15, 16] and found that body clock, sleep duration, and coffee intake impact alertness cycles.

Data from smartphones and wearable devices has extensively been used for modeling daily behavior patterns such as movement [17], sleep [18], and physical and social activities [19] to understand their associations with health and wellbeing. For example, Medan et al. [20] found that decreases in call, SMS messaging, Bluetooth-detected contacts, and location entropy (a measure of the popularity of various places) were associated with greater depression. Wang et al. [21] monitored 48 students’ behavior data for one semester and demonstrated that there are many significant correlations between the data from the smartphone and the students’ mental health and educational performance. In addition, Saeb at al [22] extracted features from GPS location and phone usage data and applied a correlation analysis for the relationship between features and level of depression. They find that circadian movement (regularity of the 24h cycle of GPS change), normalized entropy(mobility between favorite locations), location variance (GPS mobility independent of location), phone usage features, usage duration, and usage frequency were highly correlated with the depression score. Doryab et al. [23] studied loneliness detection through data mining and machine learning modeling of students’ behavior from smartphone and Fitbit data and showed different patterns of behavior related to loneliness including less time spent off campus and in different academic facilities as well less socialization during evening hours on weekdays among students with high level of loneliness.

Recent tools such as Rhythomic [24] and ARGUS [25] use visualization to analyze human behavior. Rhythomic is an open-source R framework tool for general modeling of human behavior, including circadian rhythms. ARGUS, on the other hand, focuses on visual modeling of deviations in circadian rhythms and measures their degree of irregularity. Through multiple visualization panes, the tool facilitates understanding of behavioral rhythms. This work is related to our computational framework for modeling human rhythms. However, in addition to the underlying assumption of, and a focus on, circadian rhythms only, these tools primarily enable understanding of rhythms through visualization whereas, in our framework, we provide means for processing different data sources, extracting information from them and discovering and modeling rhythms for each biobehavioral signal with different periods other than 24 hours. To our knowledge, this is the first computational framework to extract and incorporate the parameters obtained from rhythm models in a machine learning pipeline to predict different outcomes.

## CoRhythMo Framework for Modeling Biobehavioral Rhythms

Our proposed framework (Figure 1) incorporates data streams from mobile and wearable devices including behavioral signals such as movement, audio, Bluetooth, wifi, and GPS and logs of phone usage and communication (calls and messages); and biosignals such as heart rate, skin temperature, and galvanic skin response. These signals are processed, and granular features that characterize biobehavioral patterns such as activity, sleep, social communication, work, and movements are extracted. The data streams of biobehavioral sensor features are segmented into different time windows of interest and sent to a rhythm discovery component that applies periodic functions on each windowed stream of the sensor feature to detect their periodicity. The detected periods are then used to model the rhythmic function that represents the time series data stream for that sensor feature. The parameters generated by the rhythmic function are used in two ways. First, they are aggregated and further processed to characterize the stability or variation in rhythms over a certain time segment. Second, they are used as features in a machine learning pipeline to predict an outcome of interest (*e*.*g*., health status). The following sections provide details on the methods used in different components of the framework.

**Fig 1.**
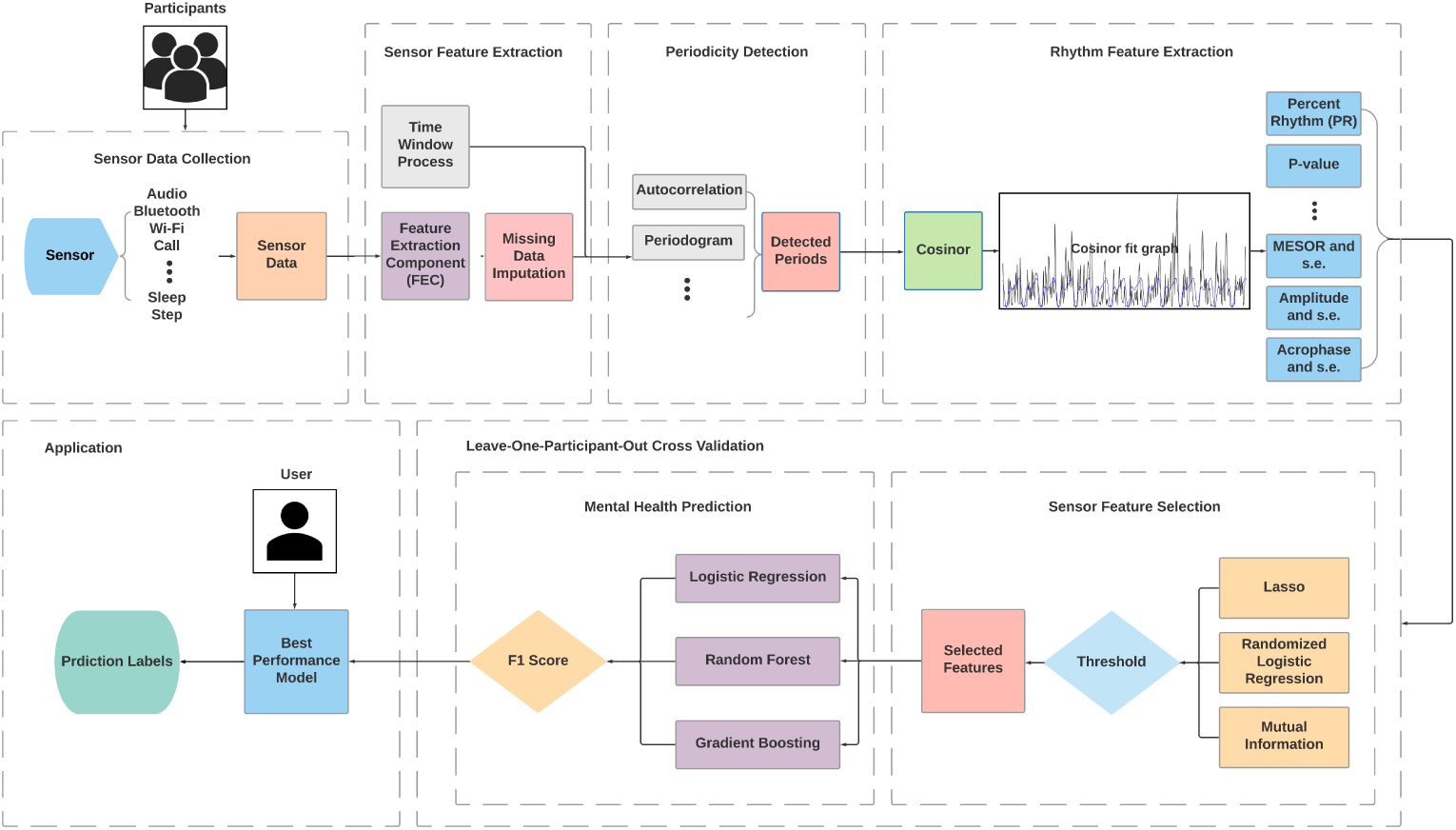
Computational framework for modeling rhythms from mobile and wearable data streams and using the rhythm parameters for prediction of an outcome (*e*.*g*., health)

### Time Series Segmentation

Windowing is one of the most frequently used processing methods for streams of data. A time series of length *L* is split into *N* segments based on certain criteria such as time. Our framework allows different ways to segment the time series, including the widely used tumbling windows, which are a series of fixed-sized, non-overlapping, and contiguous time intervals. We call each segment a time window (*tw*), which is a time series of length *l*, where *l* = *L/N*.

We also add a second segmentation layer to the time series where at each round *k* and starting point *s* (*s* = 1…*N*), we allow to combine a sequence of *k* consecutive time windows (*k* = 1…*N*) starting from time window *s* (*tw*_*s*_) to generate time series of length *k*. We call these segments time chunks (*tc*). For example, in round *k* = 1, the *tc*_11_ is a time chunk of length one and starting point of *tw*_1_ and *tc*_12_ is a time chunk of length one and starting point *tw*_2_ whereas for *k* = 3, the *tc*_32_ is a time chunk of length three and starting point of *tw*_2_. Time chunks allow flexible modeling of rhythms in different time periods over the length of the time series. Figure 2 illustrates the time segmentation process.

**Fig 2.**
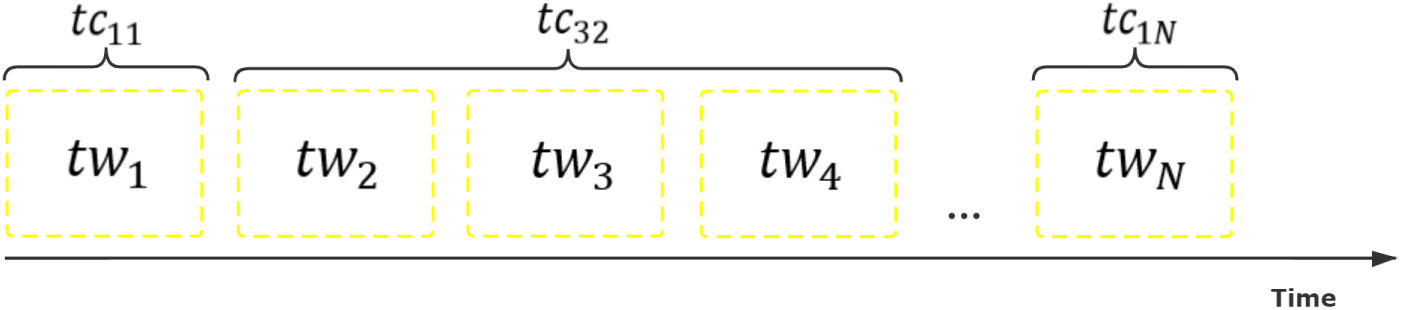
The segmentation of time series with time windows (*tw*) and time chunks (*tc*)

### Detection of Rhythmicity

One of the first steps in modeling biobehavioral rhythms is identifying rhythmicity in the time series data. We use two main methods for detecting and observing cyclic behavior, namely Autocorrelation and Periodogram.

#### Autocorrelation

Autocorrelation is a reliable analytical method for recognizing periodicities [26]. It calculates the correlation coefficient between a time series and its lagged version to measure the similarity between them over consecutive time intervals. Formally, the autocorrelation function (ACF) between two values *y*_*t*_, *y*_*t-k*_ in a time series *y*_*t*_ is defined as

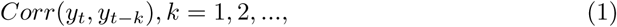

where *k* is the time gap and is called the lag [27]. In each iteration, the two time series are shifted by *k* points until one third of data is parsed. If the time series is rhythmic, the coefficient values increase and decrease in regular intervals, and significant correlations indicate strong periodicity in data. The autocorrelation sequence of a periodic signal has the same cyclic characteristics as the signal itself. Thus, autocorrelation can help verify the presence of cycles and determine the periods. It has been empirically applied on various types of time series data from different fields and was shown to be dependable and exact in the tested situations [28, 29].

#### Periodogram

A key step in the rhythm discovery process is estimation of the length of period for each rhythm. Many different techniques and algorithms for determining the period of a cycle have been developed including the Fourier-transform based methods such as Fast Fourier Transform [30], Non-Linear Least Squares [31] and Spectrum Resampling [32]. Other frequently used methods are Enright and Lomb-Scargle periodograms [33, 34], mFourfit [35], Maximum Entropy Spectral Analysis [36], and Chi-Square periodograms [37]. All of these methods come with different assumptions and with different levels of complexity [38]. For example, Spectrum Sampling has outperformed the usual Fourier approximation methods and has shown more robustness towards non-sinusoidal and noisy cycles [39]. It has also been used to detect changes in period length, which allows for estimation of variance in different periods, as frequently observed in practice. These functionalities, however, have made the algorithm slow and computationally expensive [39].

Arthur Schuster used Fourier analysis to evaluate periodicity in meteorological phenomena and introduced the term *’periodogram’* [40]. The method was first applied to the study of circadian rhythms in the early 1950s to quantify free-running rhythms of mice after blinding [41]. Periodograms provide a measure of the strength and regularity of the underlying rhythm through estimation of the spectral density of a signal. For a time series *y*_*t*_, *t* = 1, 2, …, *T*, the spectral energy *P*_*k*_ of frequency *k* can be calculated as [42]:

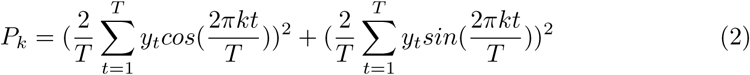

The periodogram uses a Fourier Transform to convert a signal from the time domain to the frequency domain. A Fourier analysis is a method for expressing a function as a sum of periodic components, and for recovering the time series from those components. The dominant frequency corresponds to the periodicity in the pattern.

### Modeling Rhythms

The next step in our framework is modeling the rhythmic behavior of a time series data, which is done via a periodic function. Each periodic function is among others specified by its period, average level (MESOR), oscillation degree (Amplitude), and time of oscillation optimal (Phase) [44]. The following rhythm parameters can be extracted from the model generated by the periodic function (Figure 3) [43, 45, 46]:

**Fig 3.**
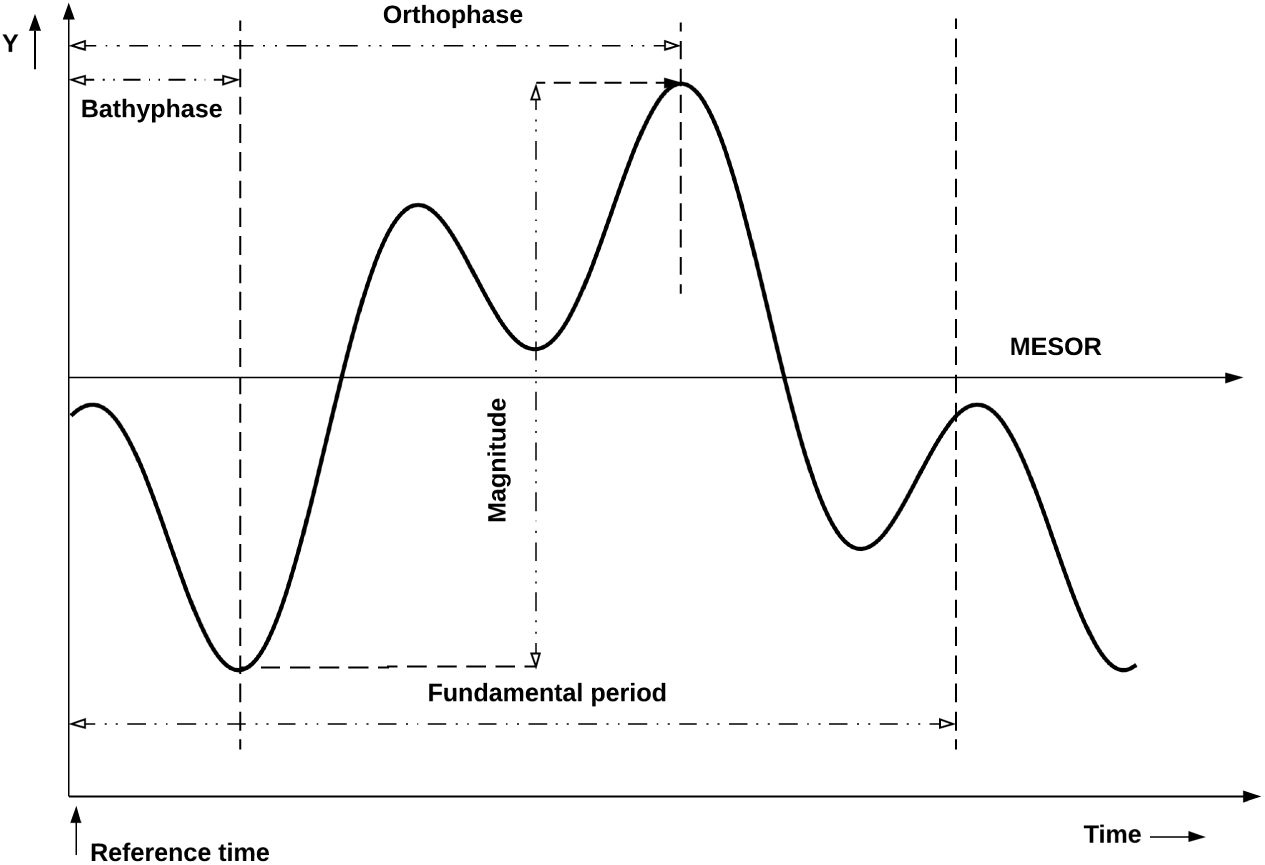
Rhythm parameters [43]

- *Fundamental period* : Periodic sequences are usually made up of multiple periodic components. The fundamental period measures the time during an overall cycle.
- *MESOR* is the midline of the oscillatory function. When the sampling interval is equal, the MESOR is equal to the mean value of all cyclic data points.
- *Amplitude* refers to the maximum value a single periodic component can reach. The amplitude of a symmetrical wave is half of its range of up and down oscillation.
- *Magnitude* refers to the difference between the maximum value and the minimum value within a fundamental period. If a periodic sequence only contains one periodic component, amplitude equals half of the magnitude.
- *Acrophase* refers to the time distance between the defined reference time point and the first time point in a cycle where the peak occurs with a period of a single periodic component.
- *Orthophase* refers to the time distance between the defined reference time point and the first time point in a cycle where the peak occurs with a fundamental period. When the time sequence only contains one periodic compoent, orthophase equals to acrophase.
- *Bathyphase* refers to the time distance between the defined reference time point and the first time point in a cycle where the trough occurs with a fundamental period.
- *P-value* indicates the overall significance of the model fitted by a single period and comes from the F-test comparing the built model with the zero-amplitude model.
- *Percent rhythm (PR)* is the equivalent to the coefficient of determination (denoted by *R*^2^) representing the proportion of overall variance accounted for by the fitted model.
- *Integrated p-value (IP)* represents the significance of the model fitted by the entire periods.
- *Integrated percent rhythm (IPR)* is the *R*^2^ of the model fitted by the entire periods.
- *The longest cycle of the model (LCM)* equals to the least common multiple of all single periods.

The most fundamental method for modeling rhythms with known periods is Cosinor, a periodic regression function first developed by Halberg et al [47] that uses the least squares method to fit one or several cosine curves with or without polynomial terms to a single time series. It uses the following cosine function to model the time series [46]:

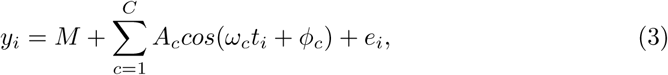

where *y*_*i*_ is the observed value at time *t*_*i*_; *M* presents the MESOR; *t*_*i*_ is the sampling time; *C* is the set of all periodic compoents; *A*_*c*_, *ω*_*c*_, *φ*_*c*_ respectively presents the amplitude, frequency, and acrophase of each periodic components; and *e*_*i*_ is the error term. In addition to the parameters described above, Cosinor outputs the standard error (SE) for MESOR, amplitude, and acrophase respectively.

The Cosinor models can be generated for one time series (single Cosinor - individual model) or for a group of time series (population-mean Cosinor - population model) through the aggregation of rhythm parameters obtained from single Cosinors. Cosinor models have been used to characterize circadian rhythms and to compute relevant parameters with their confidence limits. The model outputs the significance of the period, and it is proved that if *P ≤* 0.05, the assumed period actually exists. Our CoRhythMo framework allows for different periodic functions to be applied to the time series data using the detected periods from the previous step. We then use the rhythmic parameters measured by the Cosinor model in our machine learning pipeline, as described in the next section.

### Measuring Rhythm Stability

An important aspect of biobehavioral rhythms is their stability or deviations from normal. As mentioned previously, disruption in biological rhythms is associated with different health outcomes. As such, in our framework, we develop methods to measure the stability and variations in rhythms among individual people over different time periods (within-person) and across different population groups (between-person). Our methods employ models built by Autocorrelation and Cosinor functions to discover and measure the stability of the time series over its length. The intuition behind developing two different methods is to compare their performance in measuring rhythm stability. Besides, each method may provide unique insights that cannot be drawn from the other method.

#### Autocorrelation Sequence Stability Score (CORRES)

Recall that Autocorrelation iteratively calculates the correlation coefficient between the time series and its lagged version from start to end. The coefficient values (r) create a sequence that can be plotted by a correlogram (Figure 4) that provides a visual representation of the rhythmicity in data. The peaks above the blue dashed line in Figure 4 indicate significant correlations, and rapid decay in the amplitude of peaks indicates variation in data. To measure fluctuations in rhythms, we develop a new method to extract variability attributes from the autocorrelation sequence and further measure an overall stability score for the time series being analyzed. The process of extracting the variability parameters from the generated autocorrelation model is as follows:

**Fig 4.**
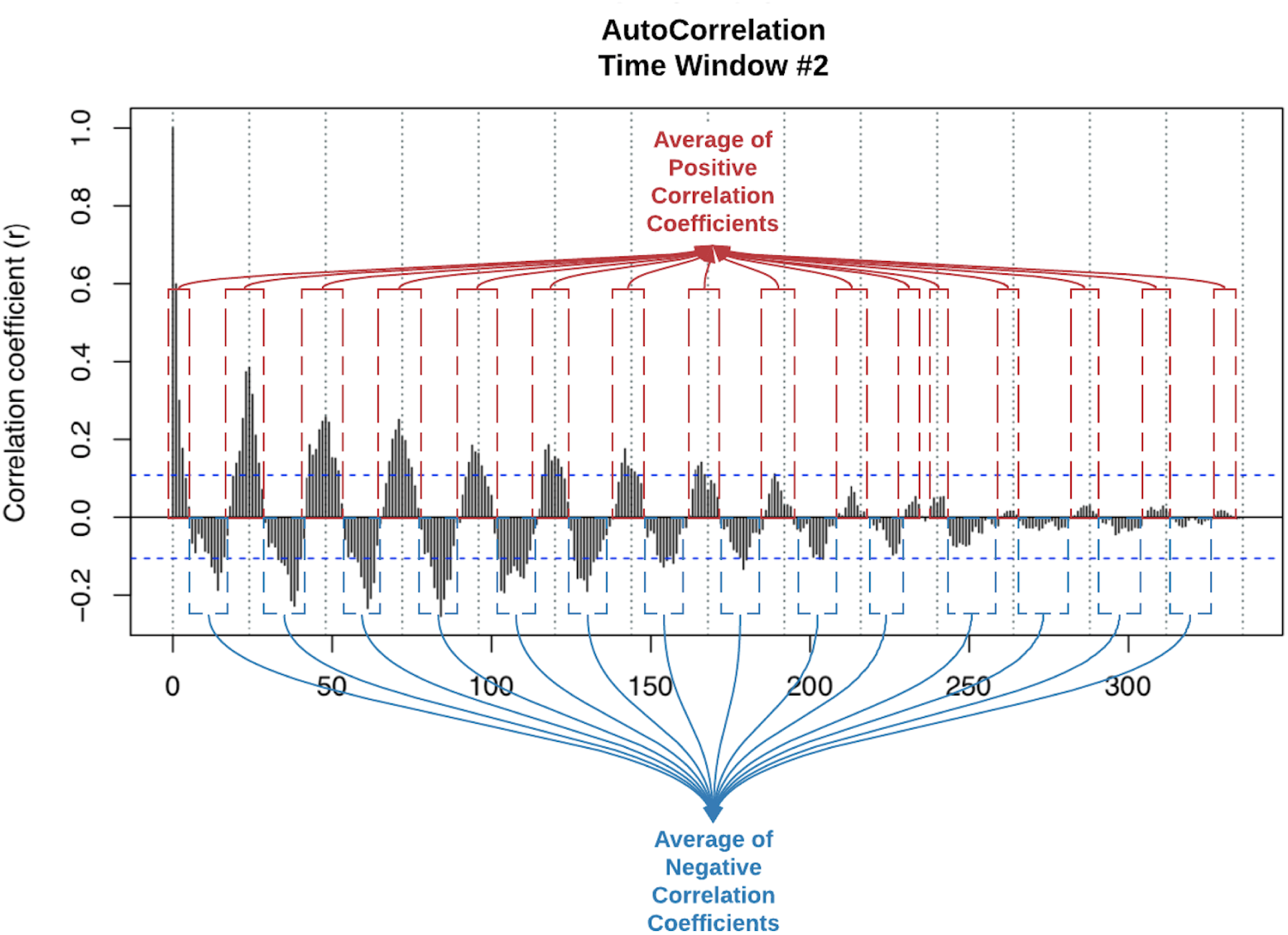
Correlogram and correlation coefficients

1. We process the generated autocorrelation sequence to calculate the mean and standard deviation of positive correlation coefficients and the mean and standard deviation of negative correlation coefficients generated by the Autocorrelation analysis as shown in Figure 4.
2. We measure the length of each correlation bout, *i*.*e*., the series of consecutive positive or negative correlations (the cones in the correlogram in Figure 4). We then calculate the min, max, mean, and standard deviation of lengths of those positive and negative bouts.
3. We identify the longest positive and negative correlation bouts in the time series and calculate the min, max, mean, and standard deviation of correlation coefficients in each of these two bouts.
4. We sort the positive correlation bouts by the highest correlation value in the bout. We then take the three positive correlation bouts with the highest correlation values and calculate the min, max, mean and standard deviation of correlation coefficients in each of the three bouts. We also calculate the same values for negative correlation bouts.

To calculate the stability, we first compute the ratio of aforementioned attributes: the positive correlation attribute over the negative correlation for that attribute (*e*.*g*., the average length of positive correlation bouts over the average length of negative correlation bouts) which provide the local stability score for that attribute. We then calculate the overall stability score by aggregating the scores of all those attributes. More formally, let *a*_*i*_ be the attribute in the above list (*e*.*g*., average correlation) and *pos*(*a*_*i*_) and *neg*(*a*_*i*_) be the corresponding positive and negative values of *a*_*i*_. The local *css css neg*(*a*) stability score (*loc*) is calculated as: 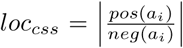. The total stability score for a *I* time window *t* is 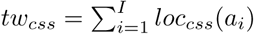 where *I* is the number of attributes. The Horizontal CORRES (HORRES) is then measured for individual time series *ts* of *K* consecutive time windows, where 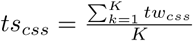. Vertical CORRES (VORRES) is measured for groups of time series (different population groups) where the CORRES score for each population group *g* of size *N*_*g*_ in each time window *tw* is measured as 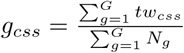 where *G* is the total number of population groups.

#### Variance of Rhythmic Parameters (COSANOVA)

As mentioned previously, Cosinor analysis can be applied to a single or group of time series. The latter is called population-mean Cosinor. We develop a second method which we call COSANOVA for measuring rhythms stability among different populations through measuring the variance of rhythm parameters obtained from the population cosinor model. COSANOVA measures the variance of MESOR, amplitude, and acrophase for each population across consecutive time windows - Horizontal COSANOVA (HANOVA), and for each time window across different populations - Vertical COSANOVA (VANOVA). The COSANOVA stability score is calculated using the average p-values from HANOVA and VANOVA. In HANOVA, the mean value and the standard error of the rhythm parameters in *K* consecutive time windows (*tw*_*t*_ - *tw*_*t*+*k*_) are used to calculate the significance (*p* value) of the variance between the means. In VANOVA, on the other hand, the mean and standard error of the rhythm parameters among *G* population groups are compared for the significance of variance in each time window *tw*_*i*_. In other words, HANOVA defines group level stability, and VANOVA defines time window level stability. If HANOVA score is greater than the significance level (*e*.*g*., 0.05), the group rhythm is stable. Similarly, if VANOVA score is greater than the significance level (*e*.*g*., 0.05), the time window rhythm is stable. The scores greater than the significance level mean the variance of rhythm is not significant.

Let *p*_*r*_ be the significance of variance for each rhythm parameter across *K* time windows for population *g*. The HANOVA score for this population is calculated as 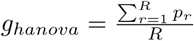 where *R* is the number of rhythm parameters. The VANOVA score for time window *tw*_*k*_ is measured as 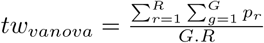 where *G* is the number of population groups. The COSANOVA score for each sensor feature *f* is the 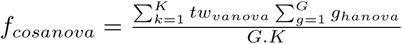

We then calculate the percentage of sensor features with stable rhythms in each population group and across time windows, which provides an overall stability score for the entire population (*i*.*e*., all groups together). Figure 5 illustrates the pipeline of calculating the stability score for the rhythms of each sensor feature.

**Fig 5.**
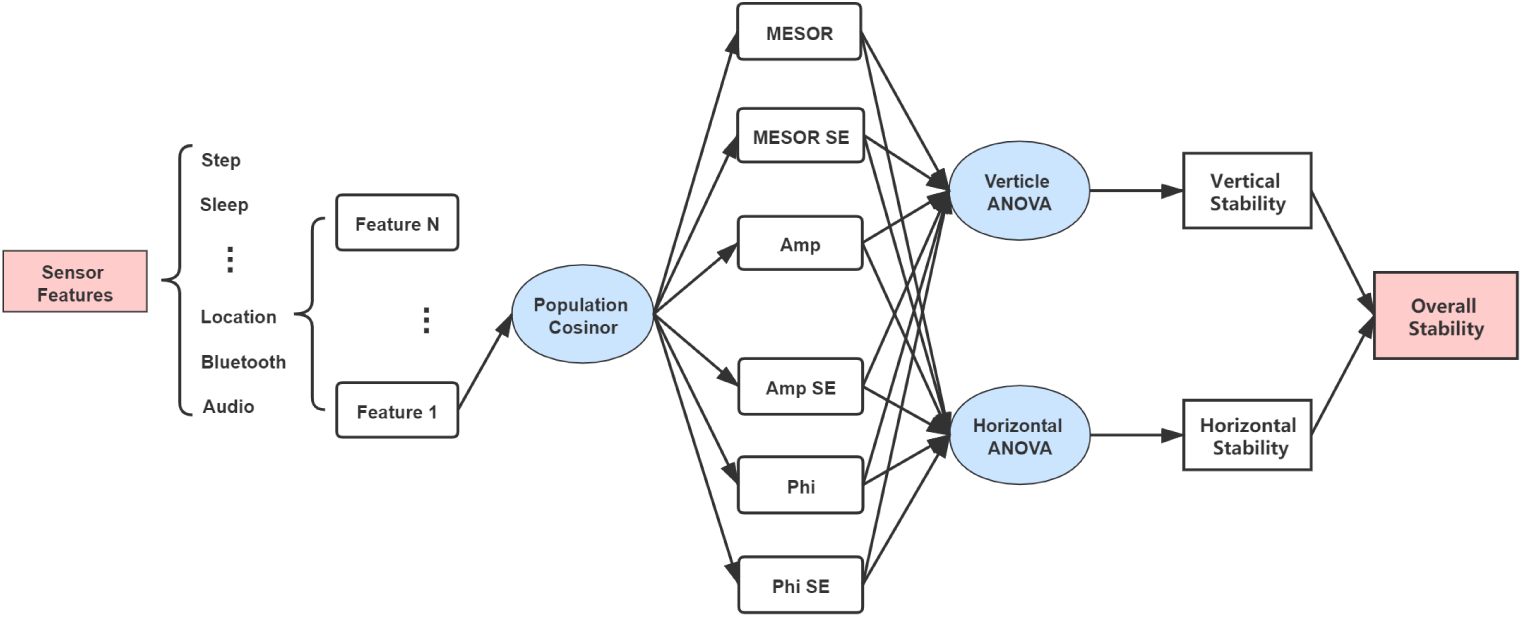
The process for measuring rhythm stability parameters

### Machine Learning Method

The machine learning component of the framework uses the parameters obtained from modeling the rhythm of each sensor feature to generate datasets for training and testing of an outcome of interest, *e*.*g*., health. The pipeline processes and handles missing values both in sensor and rhythm features across different time windows select important rhythm features as part of the training process and builds machine learning models for prediction of the outcome. The following sections describe the details of each step.

#### Handling Missing Values

Given the streams of data from multiple sources, the framework handles missing data for each sensor stream and each time window. We remove any sensor feature if the percent of its missing data is greater than a threshold (*e*.*g*., 30%). For remaining sensor features, we perform nearest-neighbor linear interpolation [48] to fill in missing values. For example, if there are three missing data points between 10 and 50, then the three missing points are filled with 20, 30, and 40, respectively. Given that the first and last data points cannot be imputed using this method, we remove the sensor feature if the first or the last data point in the time window is missing.

We apply the same process for handling missing rhythmic features in consecutive time windows. For each rhythmic feature, we fill the value of the missing time window with nearest-neighbor linear interpolation. Let *v*_*i*_ be the value of time window *tw*_*i*_. If *v*_1_ and *v*_5_, the values of time windows *tw*_1_ and *tw*_5_ are present and *v*_2_, *v*_3_, and *v*_4_, the *-*values of *tw*_2_, *tw*_3_ and *tw*_4_ are missing, then 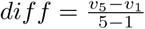, and *v*_2_ = *v*_1_ + *diff, v*_3_ = *v*_1_+ *diff ∗* 2, and *v*_4_ = *v*_1_ + *diff ∗* 3. For each missing time window, if none of the time windows before it has value, or none of the time windows after it has value, then this time window is not filled. After imputation, we remove any rhythmic feature with missing values more than a threshold (*e*.*g*., 30%). Algorithm 1 describes the process in more details.

##### Algorithm 1: Missing value imputation

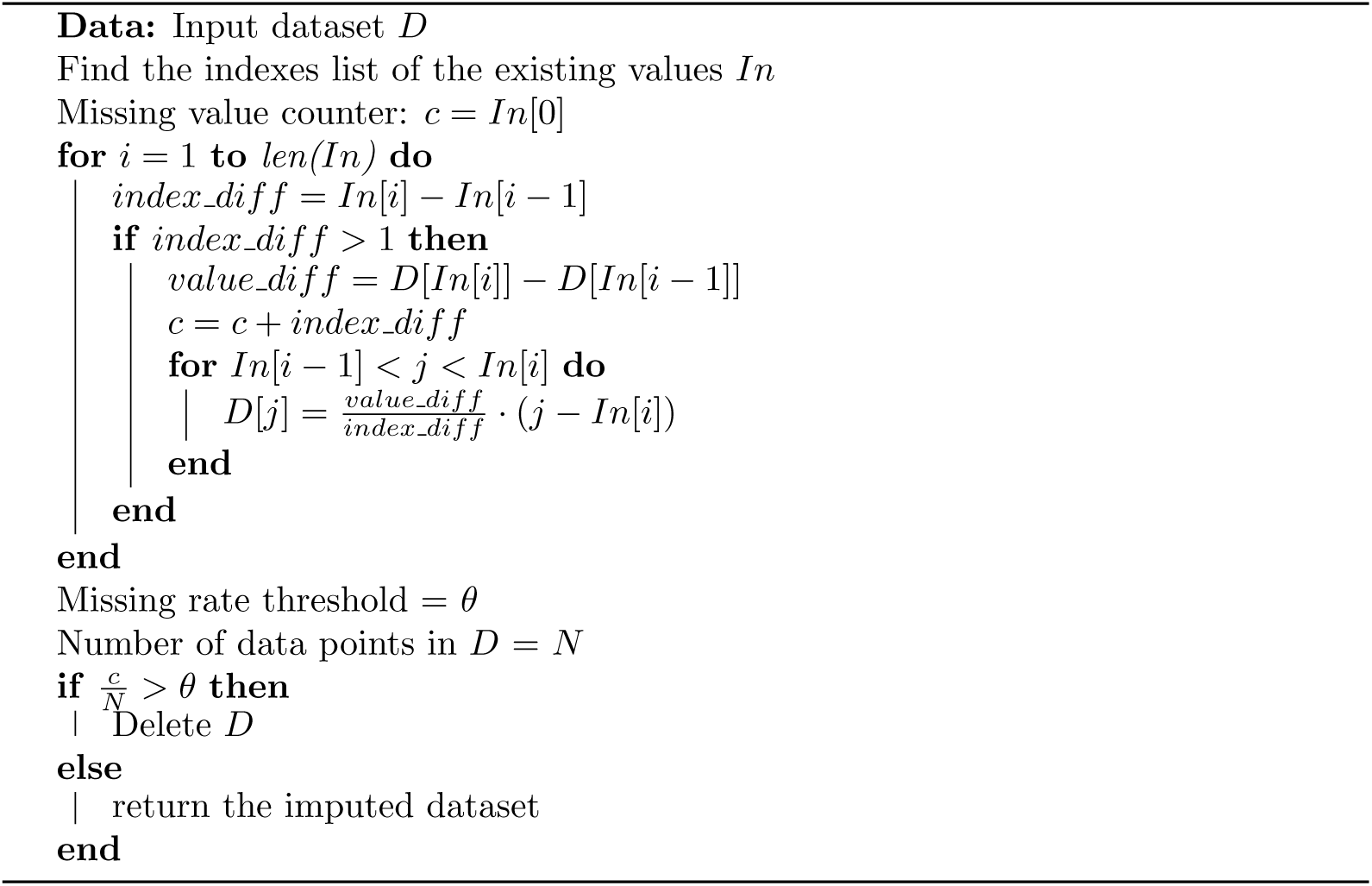

#### Feature Selection

As mentioned in previous sections, for each type of sensor feature, a single period or a multi-frequency Consinor model is generated, which outputs a list of rhythm parameters. These parameters are entered the training process for building machine learning models.

Let *M* be the number of sensors (*s*_1_…*s*_*m*_), *FN*_*i*_ be the number of features for sensor *i* and *RN*_*j*_ the corresponding number of rhythmic features for feature *j* in sensor *i*. The resulting feature space will be of *M ∗ FN ∗ RN*, which is high dimensional compared to the relatively few data samples for training. As such, a reduction in the number of features is prevalent. The framework allows for the integration of different feature selection methods. To start, we implement three widely used methods, namely Lasso, Randomized Logistic Regression (RLR), and Information Gain(IG), in the machine learning component.

*Lasso* is a linear regression model penalized with the L1 norm to fit the coefficients [49]. The Lasso regression prefers solutions with fewer non-zero coefficients and effectively reduces the number of features that are independent of the target variable. Through cross-validation, the lasso regression can output the importance level for each feature in the training dataset. We use a threshold value of 1e-5 to select features with Lasso, which is the default threshold in Sklearn library. Features with importance greater or equal to the threshold are kept, and the rest are discarded.

*Randomized Logistic Regression* is developed for stability selection of features. The basic idea behind stability selection is to use a base feature selection algorithm like logistic regression to find out which features are important in bootstrap samples of the original dataset [50]. The results on each bootstrap sample are then aggregated to compute a stability score for each feature in the data. Features with a higher stability score than a threshold are selected. We use 0.25, the default threshold value in Sklearn library. Note this stability score is for feature selection and is different from stability measures we developed for rhythms.

*Information Gain* (also referred to as Mutual Information in feature selection) measures the dependence between the features and the dependent variable (predicted outcome) [51]. Mutual information is always larger than or equal to zero, where the larger the value, the greater the relationship between the two variables. If the calculated result is zero, then the variables are independent. We set our algorithm to select 10 (the default value in Sklearn library) features with the highest information gain.

#### Model Building and Validation

The step for building machine learning models using rhythm features of *k* consecutive time windows and for a population of *D* data samples is flexible in the framework and can incorporate different supervised and unsupervised machine learning methods such as regression, classification, and clustering. In the current version of the framework, we implement three classification methods, including *Logistic Regression (LR), Random Forest (RF), and Gradient Boosting (GB)*. The choice of algorithms is simply based on our empirical evidence of their performance on this type of data. Logistic regression [52] uses the logistic function to build a classifier. Random forest and Gradient Boosting are two branches of ensemble learning [53], which use the idea of bagging and boosting [54] respectively. Their common feature is to use the decision tree as the basic classifier and to get a robust model by combining multiple weak models. Bagging is short for boost strapped aggregation. Boost strapping is a repeated sampling method with putting back, and the sampling strategy is simple random sampling [55]. In boosting, classifiers are connected in series, where the training set of each round is unchanged, but the weight of samples is changed. At each iteration, the training samples with high error rates are given higher weights, so they get more attention in the next round of the classifier.

In order to better understand the role of each sensor in prediction, we build models with features from single sensors alone and features from multiple sensors. We use a baseline of the majority class to measure the performance of the classifiers in the prediction of the outcome. Again, the flexibility of the framework allows for the incorporation of different baseline measures. Both feature selection process and building machine learning models are done in a cross-validation setting, *e*.*g*., leave one sample out [56]. The machine learning component can measure basic performance measures of accuracy, precision, recall, F1, and MCC scores to evaluate the performance of the algorithms. From those measures, we choose the results above baseline for each combination of feature selection and learning algorithm to further explore the prediction outcomes and to gain insights.

### Evaluation

We utilized a dataset of smartphone, Fitbit, and survey data collected from 138 first-year undergraduate students at an American university who were recruited for a health and well-being research study. The dataset was previously used in [23] to detect loneliness among college students. Smartphone data was collected through the AWARE framework [57] and included calls, messages, screen usage, Bluetooth, Wi-Fi, audio, and location. A Fitbit Flex2 wearable fitness tracker tracked steps, distances, calories burned, and sleep; and survey questions gathered information about physical and mental health, including loneliness and depression. The survey data was collected at the beginning and at the end of the semester.

### Data Processing

#### Survey Data

In our evaluation, we focused on two mental health outcomes namely depression and loneliness. These two measures were chosen because of their longitudinal aspect, *i*.*e*., lasting for at least a few weeks to enable the investigation of 1) how biobehavioral rhythms of students with mental health conditions would differ from other students and how well the state of those mental health conditions could be predicted from extracted rhythms.

Loneliness data was collected using the UCLA Loneliness Scale, a well-validated and commonly used measure of general feelings of loneliness [58]. The questionnaire contains 20 questions about feeling lonely and isolated using a scale of 1 (never) to 4 (always). The total loneliness scores range from 20 to 80, with higher scores indicating higher levels of loneliness. As there is no standard cutoff for loneliness scores in the literature, we followed the same approach in [23] to divide the UCLA scores into two categories where the scores of 40 and below were categorized as *’low loneliness’*, and the scores above 40 were categorized as *’high loneliness’*.

Depression was assessed using the Beck Depression Inventory-II (BDI-II) [59, 60], a widely used psychometric test for measuring the severity of depressive symptoms that have been validated for college students [60]. The BDI-II contains 21 questions, with each answer being scored on a scale of 0-3 where higher scores indicate more severe depressive symptoms. For college students, the cut-offs on this scale are 0-13 (no or minimal depression), 14-19 (mild depression), 20-28 (moderate depression) and 29-63 (severe depression) [60]. For simplicity and to be consistent with the loneliness categorization, we divided these scores into two categories where the BDI-II scores *<* 14 were labeled as *’not having depression’*, and all BDI-II scores *>*= 14 were labeled as *’having depression’*.

These loneliness and depression categories were used as ground truth labels in our machine learning pipeline to classify students’ depression and loneliness levels using rhythmic features. Since there are two surveys for each student at the beginning and at the end of the semester, students could be separated into five groups according to the mental health surveys for depression and loneliness. For simplicity of representation, we further label *low loneliness* and *no depression* categories as 1, and *high loneliness* and *high depression* as 2. The five mental health categories are as follows:

- All students
- Pre1 Post1: not having a mental health condition in both pre-semester and post-semester surveys
- Pre1 Post2: not having a mental health condition in the pre-semester survey, but having it in the post-semester survey
- Pre2 Post2: having a mental health condition in both surveys
- Pre2 Post1: having a mental health condition in the pre-semester survey, but not in the post-semester survey

#### Sensor Data

The dataset collected from smartphones and Fitbits consisted of time series data output by multiple sensors, including Bluetooth, calls, SMS, Wi-Fi, location, phone usage, steps, and sleep. We grouped this time series data into hourly bins and processed it following the approach in [61] to extract features related to mobility and activity patterns, communication and social interaction, and sleep. Examples of such features include travel distance, sleep efficiency, and movement intensity. We then split the semester data into tumbling cyclic time windows of 14 days or two weeks based on empirical evaluation of different lengths of time windows. The university semester in the studied population was roughly 16 weeks long, which could be divided into eight time windows of two weeks except for the last time window that contained only ten days of data (Figure 6). We built a model of rhythm for each student and for each time window.

**Fig 6.**
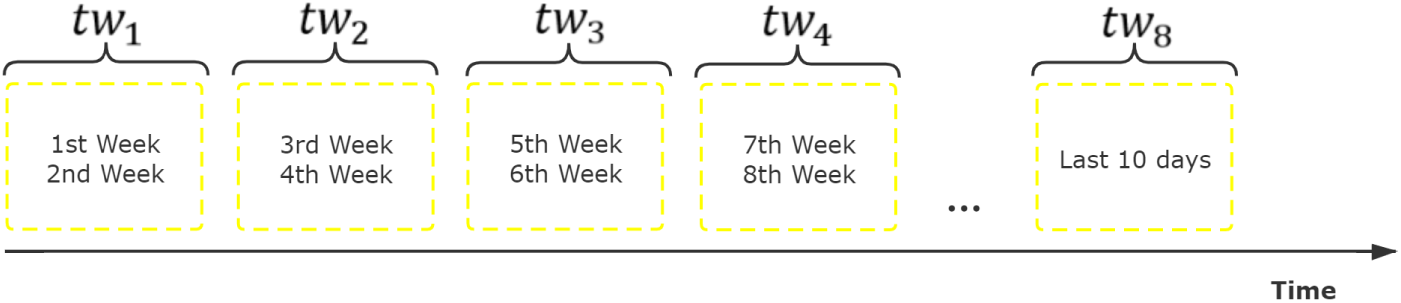
The size of a time window is 2 weeks which segments the semester into roughly 8 time windows.

We handled missing sensor data on a per-participant per-time window basis. For each participant and each time window, we removed any sensor feature if it had more than 30% missing data. For remaining sensor features, we performed nearest-neighbor linear interpolation as described previously to fill in missing values.

#### Analysis and Results

Our analysis was performed in two steps: First, we explored the potential of modeling and detecting rhythmicity in passively collected data from students’ mobile and wearable data streams. Then we used the built rhythm models to extract features that were fed into machine learning models to explore the relationship between students’ biobehavioral rhythms and their mental health. We aimed to answer the following questions:

- Can we observe rhythmicity in students’ data over the course of the semester? If so, are those rhythms consistent throughout the semester, or do they change during different periods?
- Do we observe any difference in biobehavioral rhythms among students with different health statuses? If so, do healthy students have more stable rhythms?
- Can models of biobehavioral rhythms predict mental health status?
- What are the most important characteristics and rhythmic features that reveal changes in health status?

The following sections describe our observations and findings. To distinguish the mental health groups in the two conditions, we add an *L* and *D* to the mental health group for loneliness (*e*.*g*., L_Pre1_Post) and depression (*e*.*g*., D_Pre1_Post) respectively.

#### Detection of rhythmicity in student data

To investigate whether we can observe rhythmicity in data collected from students’ smartphones and Fitbits, we used Autocorrelation, Periodogram, and Cosinor to model students’ rhythms in each time window.

Our results shows that the most dominant periods in each time window are 24- and 12-hours. As shown in Tables 1 and 2, for sleep duration feature in depression category, this trend is consistent in all students regardless of the mental health condition where 97.6% and 69.6% of students have 24- and 12-hours as dominant periods in their data. The 12-hour period has the lowest percentage of students (46.3%) in group D_Pre2_Post2 which includes students who were depressed throughout the semester).

**Table 1.**
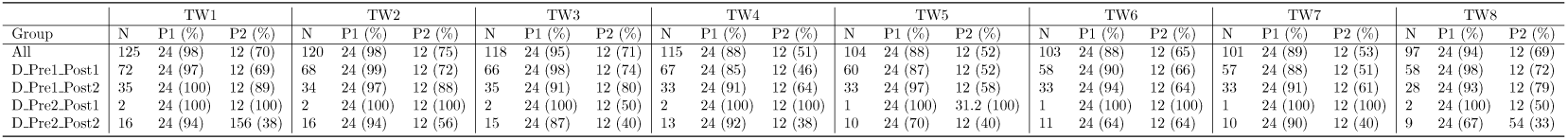
Top two dominant periods of num asleep feature for depression groups. N is the number of students in the group. P1 is the most dominant period (*i*.*e*., the percentage of students that have the period is highest among all periods). The percentage in parenthesis is the percentage of students that have the period. P2 is the second dominant period.

**Table 2.** Top three dominant periods of num asleep (minutes asleep) feature for Pre1 Post2 groups. N is the number of students in the group. P1 is the most dominant period (*i*.*e*., the percentage of students that have this period is highest among all periods). The percentage in parenthesis is the percentage of students that have the period. P2 is the second dominant period and P3 is the third dominant period.

| Pre1_Post2 |  |  |  |  |  |  |  |  |
| --- | --- | --- | --- | --- | --- | --- | --- | --- |
| Time Window | Loneliness |  |  |  | Depression |  |  |  |
|  | N | P1 (%) | P2 (%) | P3 (%) | N | P1 (%) | P2 (%) | P3 (%) |
| TW1 | 17 | 24 (100) | 12 (71) | 312 (35) | 35 | 24 (100) | 12 (89) | 312 (34) |
| TW2 | 15 | 24 (93) | 12 (87) | 312 (40) | 34 | 24 (97) | 12 (88) | 312 (38) |
| TW3 | 16 | 24 (100) | 12 (88) | 156 (31) | 35 | 24 (91) | 12 (80) | 156 (31) |
| TW4 | 15 | 24 (73) | 12 (53) | 312 (33) | 33 | 24 (91) | 12 (64) | 78 (40) |
| TW5 | 14 | 24 (100) | 12 (64) | 156 (29) | 33 | 24 (97) | 12 (58) | 312 (36) |
| TW6 | 12 | 24 (92) | 12 (67) | 78 (33) | 33 | 24 (94) | 12 (64) | 78 (45) |
| TW7 | 13 | 24 (85) | 12 (54) | 156 (31) | 33 | 24 (91) | 12 (61) | 156 (40) |
| TW8 | 11 | 24 (91) | 12 (55) | 72 (45) | 28 | 24 (93) | 12 (78) | 72 (32) |

Table 2 shows that the dominant periods of 24- and 12-hours are preserved for the sleep duration feature in all time windows for students with loneliness and depression even though students in both conditions started the semester with normal health status but developed depression or loneliness towards the end (D Pre1 Post2). The lowest percentage of students in this group with 24- and 12-hour periods are in time windows 4 and 5 with 73% in loneliness category (24-hour), 91% in depression category (24-hour), 53% in loneliness category (12-hour), and 57% in depression category (12-hour).

Overall and across all sensor features, we observe the 24-hour as the dominant period for over 52% of the student population with the highest percentages belonging to steps (95%), calories (92%), wifi (83%), and sleep (68%). Table 3 presents the overall percentages for each sensor. Following these observations, we looked at the percentage of participants in each mental health group that had 24-hours as one of their dominant rhythms for each *time chunk*. This would help observe the extent to which students preserved their normal circadian rhythm over the semester. Recall that time chunks consist of *k* consecutive time windows, there were 36 different time chunks in total for eight time windows of length 2 in the dataset. In each time chunk, a participant had 24-hour as a dominant rhythm if and only if this participant had 24-hour as a dominant rhythm in all time windows in that time chunk. Figure 7 shows the percentage of participants with 24-hour as the dominant rhythm (y-axis) in each mental health group for each time chunk of length 3 (x-axis). The data point at *x* = *i* corresponds to the time chunk of length 3 starting at *tw*_*i*_ (*i*.*e*., *tc*_3*i*_). It represents the percentage of participants with 24-hour as the dominant rhythm in all the 3 time windows *tw*_*i*_, *tw*_*i*+1_, *tw*_*i*+2_ (6 consecutive weeks).

**Table 3.** Percentage of participants with 24-hour period across all sensor features

| % of Participants with 24-hour period |  |  |  |  |  |  |  |  |  |  |
| --- | --- | --- | --- | --- | --- | --- | --- | --- | --- | --- |
| Audio | Battery | Bluetooth | Calorie | Location | Location Map | Call&Messages | Screen | Sleep | Steps | Wifi |
| 62 | 13 | 42 | 92 | 41 | 17 | 18 | 36 | 69 | 95 | 83 |

**Fig 7.**
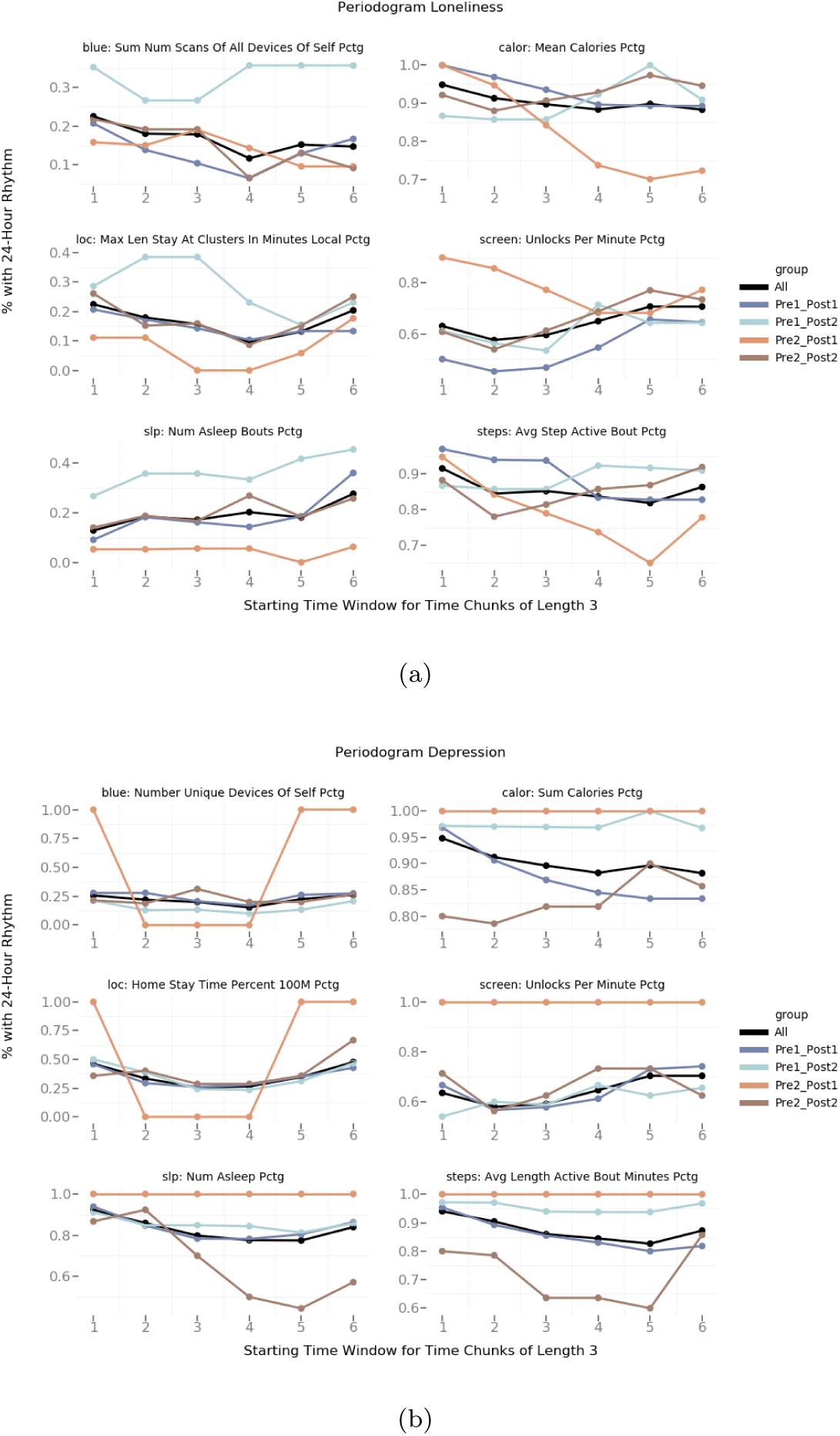
The plots show the percentage of participants with 24-hour as the dominant rhythm (y-axis) in each mental health group (left: loneliness, right: depression) for each time chunk of length 3 (x-axis). The data point at *x* = *i* corresponds to the time chunk of length 3 starting at *tw*_*i*_ (*i*.*e*., *tc*_3*i*_). It represents the percentage of participants with 24-hour as the dominant rhythm in all the 3 time windows *tw*_*i*_, *tw*_*i*+1_, *tw*_*i*+2_.

For loneliness, the group with low loneliness at the beginning and high loneliness at the end of the semester (L_Pre1_Post2) has a higher percentage of 24-hour rhythms for features of sleep, location, and Bluetooth. The opposite group with high loneliness at the beginning and low loneliness at the end of the semester (L_Pre2_Post1) has a lower percentage of 24-hour rhythms for features of calories and steps, while it has higher percentage for screen features. For depression, the group D_Pre2_Post1 included only one participant in that group, while other groups had more than ten people. The group with no depression at the beginning and with depression at the end of the semester (D_Pre1_Post2) had a higher percentage of normal 24-hour rhythms for features of calories and steps. The group that was depressed throughout the semester (D_Pre2_Post2) had a lower percentage of 24-hour rhythms for steps, sleep, and calories. All groups except D_Pre2_Post1 had a similar percentage for Bluetooth, location, and screen.

#### Measuring the stability of rhythms

The second analysis question concerned the stability of rhythms among different mental health groups. We used Autocorrelation Sequence Stability Score (CORRES) and Cosinor Anova (COSANOVA) methods we developed and described earlier for measuring the stability of the rhythms.

Figure 9 shows a visual representation of fluctuations in the correlogram attribute, which calculated the average positive and negative correlations (figure 4) observed through the autocorrelation analysis. The plots show the autocorrelation coefficient average (y-axis) of the five groups for the eight time windows (x-axis). Each plot corresponds to one sensor feature. For loneliness, the group with low loneliness at the beginning and high loneliness at the end of the semester (L_Pre1_Post2) has more variation of autocorrelation coefficient across time for the features of call, phone usage (screen) and sleep. Meanwhile, the negative autocorrelation coefficient of this group becomes lower than other groups towards the end of the semester for the features of calories, location, and steps. For depression, the two groups that are not depressed at the beginning of the semester (D_Pre1_Post1, D_Pre1_Post2) have similar autocorrelation average, while the two groups that are depressed at the beginning of the semester (D_Pre2_Post1, D_Pre2_Post2) have different autocorrelation pattern from other groups.

**Fig 8.**
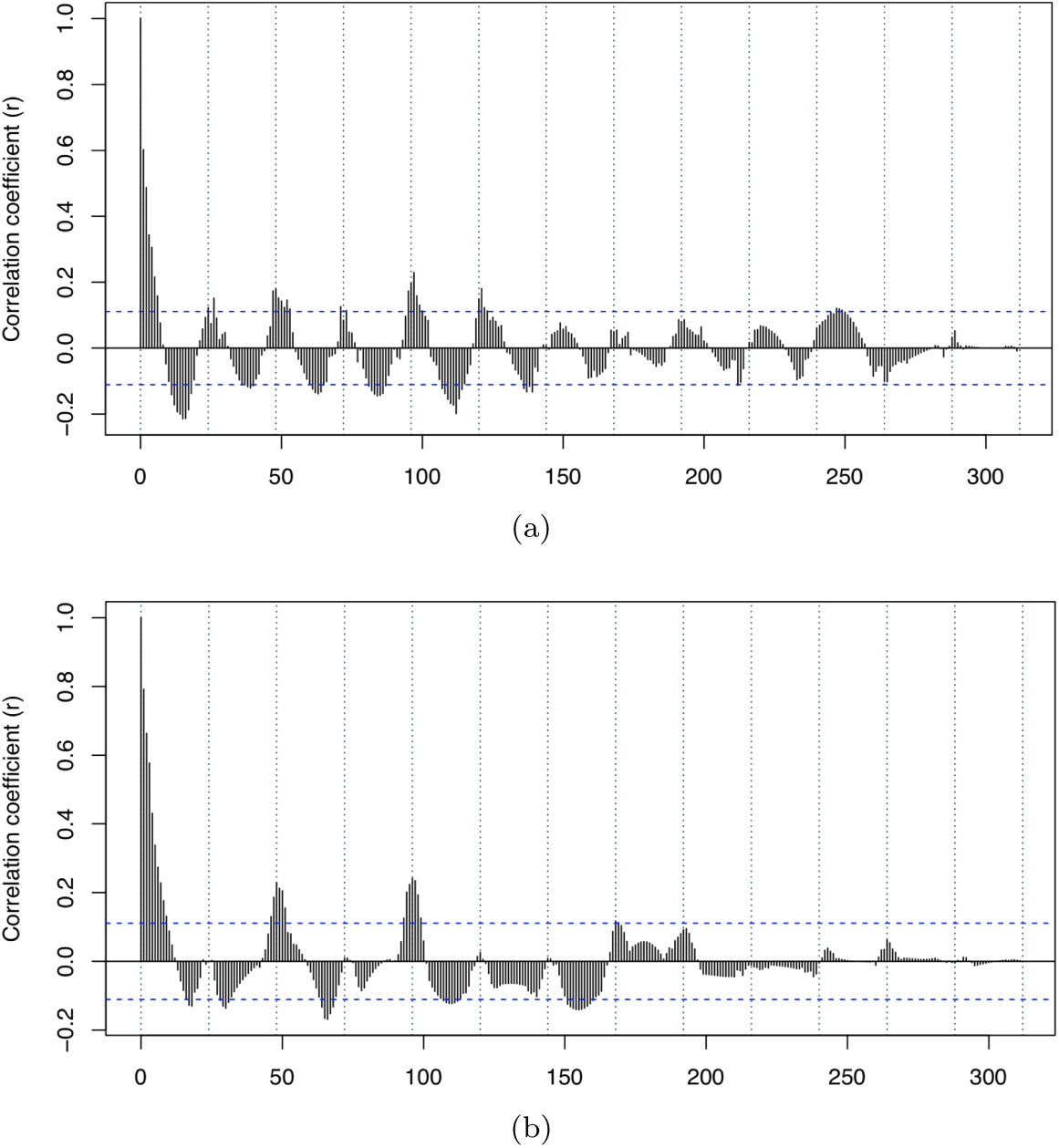
Correlograms of feature num_restless bout (number of restless periods in sleep) in time window 4 for two students (left: a student in L_Pre1_Post1, right: a student in L_Pre1_Post2).

**Fig 9.**
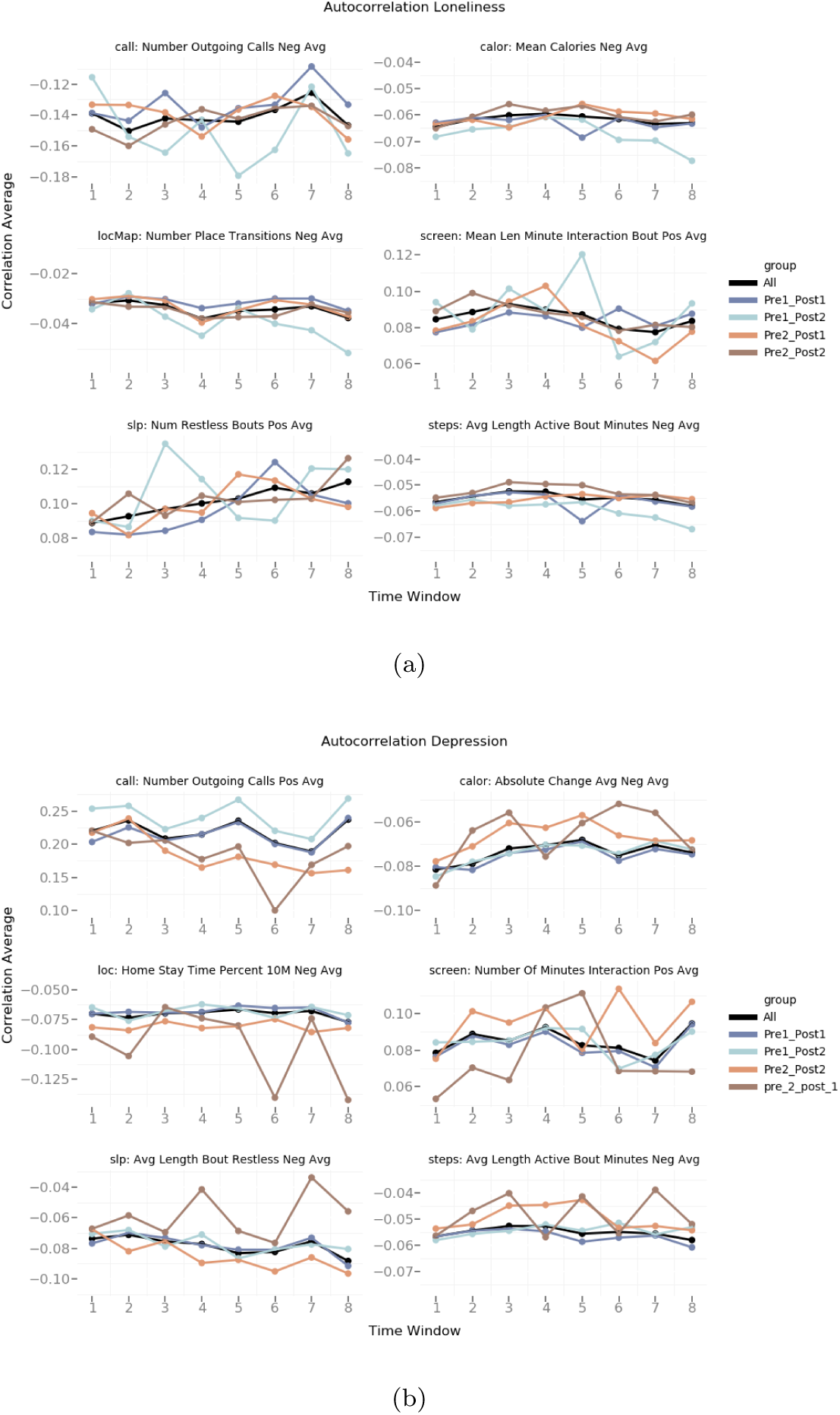
The plots show the autocorrelation coefficient average (y-axis) of the 5 groups (left: loneliness, right: depression) for the 8 time windows (x-axis). Each plot corresponds to one sensor feature.

We then applied the CORRES method to obtain the stability scores of each population group in each time window and across all time windows. As shown in Table 4, the student population as a whole shows most stability in rhythms of restless sleep in time window 1 and 7 and among the mental health groups, the L_Pre1_Post1 (students with no loneliness throughout the semester) has the most stable rhythms overall and in the majority of time windows including *tw*_1_, *tw*_2_, *tw*_5_, and *tw*_7_. Figure 8 shows the correlogram of the number of restless sleep bouts in two students from different groups, one with low loneliness throughout the semester and the other with high loneliness at the end of the semester. The figure visually depicts differences in the rhythms of these two students, where the correlogram belonging to students with high loneliness projects a less stable rhythm towards the end of time series.

**Table 4.** Stability score of Autocorrelation coefficient of feature num restless bout (number of restless periods in sleep) for loneliness groups.

| Group | TW1 | TW2 | TW3 | TW4 | TW5 | TW6 | TW7 | TW8 | H-CORRES |
| --- | --- | --- | --- | --- | --- | --- | --- | --- | --- |
| all | 115.40 | 82.73 | 60.70 | 60.08 | 94.04 | 64.71 | 134.98 | 63.12 | 84.47 |
| Pre_1_Post_1 | 261.54 | 153.62 | 64.66 | 57.75 | 173.74 | 44.00 | 312.41 | 43.61 | **138.92** |
| Pre_1_Post_2 | 52.67 | 47.00 | 53.44 | 63.67 | 57.58 | 83.21 | 70.75 | 46.22 | 59.32 |
| Pre_2_Post_1 | 76.73 | 55.31 | 55.05 | 71.16 | 63.65 | 44.39 | 55.29 | 63.85 | 60.68 |
| Pre_2_Post_2 | 54.48 | 59.20 | 62.40 | 56.47 | 61.83 | 80.52 | 50.26 | 85.60 | 63.84 |
| V-CORRES | **112.16** | 79.57 | 59.25 | 61.83 | 90.17 | 63.37 | **124.74** | 60.48 | **81.45** |

##### COSANOVA

To measure population rhythms’ stability, we first build population-mean Cosinor models of different mental health groups. We then use the values of MESOR, amplitude, and phase and their corresponding standard error (SE) provided by the Cosionr analysis to compute the HANOVA and VANOVA scores for each sensor feature, each time window, and each mental health group.

To evaluate the performance of our COSANOVA method, we first explored the Cosinor models of four different features and the fluctuation of rhythm parameters in different mental health groups and across time windows (Figure 10). We observed different oscillation patterns of rhythm features in the four mental health groups. For example, the pattern of change in the magnitude of rhythm for the feature slp max length bout restless (top left figure) that measures the maximum length of restless sleep is minor for groups L_Pre1_Post1 and L_Pre2_Post2 (*i*.*e*., groups with unchanged mental state over the semester), but the change is significant for groups L_Pre1_Post2 and L_Pre2_Post1 (*i*.*e*., groups that experience change in their mental state over the semester). In particular, while L_Pre1_Post2 has the highest magnitude (double value of amplitude) in time window 3, the L_Pre2_Post1 group shows the lowest amplitude. In contrast, the MESOR of rhythms for sum calories (the total used calories) feature shows a similar fluctuation pattern across all groups and time windows (top right figure). For example, although L_Pre1_Post1 and L_Pre2_Post2 groups have the exact opposite mental health status, their wave patterns are similar in most time windows (*e*.*g*., *tw*_6_). The bottom two figures demonstrate other cases where the same mental health group projects, similar or different rhythms in different time windows. Following these observations, we calculate the HANOVA, VANOVA, and COSANOVA for the same features to see whether the stability measurements match our observations. Table 5 illustrates that our defined HANOVA and VANOVA could detect unstable rhythm, as discussed above. For example, in the top left figure, the HANOVA for group L_Pre1_Post1 and L_Pre2_Post2 is insignificant, indicating their stability across time windows, whereas VANOVA is significant in *tw*_2_, *tw*_3_, *tw*_4_, and *tw*_5_ indicating fluctuations across mental health groups in those time windows. We repeat this process and calculate the COSANOVA score for all sensor features. As shown in Table 5, the loneliness group Pre1_Post1 (the group with low loneliness over the course of the semester) has the highest percentage of sensor features with stable rhythms (54.1%) followed by Pre2_Post2 (47.4%). The percentage of features with stable rhythms in each time window remains below 40% with *tw*_3_ as the most stable (38.3%) followed by *tw*_5_ and *tw*_6_.

**Table 5.** HANOVA and VANOVA results of sensor features including features shown in Figure 10. The values in the table are the p-values outputted by the two ANOVAs. Percentages represent the proportion of stable features to the total number of features in each group or each time window.

| HANOVA |  |  |  |  |  |  |  |  |  |
| --- | --- | --- | --- | --- | --- | --- | --- | --- | --- |
| Features | Pre1_Post1 | Pre1_Post2 | Pre2_Post1 | Pre2_Post2 | H-COSANOVA |  |  |  |  |
| slp_max_length_bout_restless | 0.3857 | 3.48E-35 | 4.21E-09 | 0.3322 | 0.1795 |  |  |  |  |
| slp_start_time_min_bout_totalsleep | 0.1324 | 0.1976 | 0.8252 | 0.5833 | 0.4346 |  |  |  |  |
| steps_avg_step_active_bout | 0.1046 | 0.06 | 0.0008 | 8.89E-05 | 0.0413 |  |  |  |  |
| calor_sum_calories | 7.58E-16 | 1.80E-11 | 2.30E-13 | 2.36E-25 | 4.56E-12 |  |  |  |  |
| blue_avg_num_scans_of_all_devices_of_others | 0.5282 | 0.3252 | 0.0006 | 5.55E-05 | 0.2135 |  |  |  |  |
| . | . | . | . | . | . |  |  |  |  |
| . | . | . | . | . | . |  |  |  |  |
| Proportion of stable sensor features per group | 54.12% | 45.88% | 45.83% | 47.43% | 42.56% |  |  |  |  |
| VANOVA |  |  |  |  |  |  |  |  |  |
| Features | TW 0 | TW 1 | TW 2 | TW 3 | TW 4 | TW 5 | TW 6 | TW 7 | V-COSANOVA |
| slp_max_length_bout_restless | 0.0203 | 0.0033 | 2.18E-16 | 4.03E-58 | 1.34E-09 | 2.28E-30 | 0.1972 | 0.0096 | 0.0288 |
| slp_start_time_min_bout_totalsleep | 0.8163 | 0.4734 | 0.0321 | 0.8783 | 0.8173 | 0.9253 | 0.499 | 0.0906 | 0.5665 |
| steps_avg_step_active_bout | 9.74E-08 | 5.96E-08 | 1.55E-10 | 3.63E-09 | 2.17E-05 | 1.51E-08 | 1.10E-09 | 2.06E-10 | 2.73E-06 |
| calor_sum_calories | 5.29E-13 | 3.23E-08 | 1.61E-09 | 8.19E-16 | 1.12E-17 | 2.60E-18 | 1.44E-17 | 4.26E-14 | 4.24E-09 |
| blue_avg_num_scans_of_all_devices_of_others | 2.64E-07 | 1.36E-27 | 1.13E-09 | 6.12E-22 | 4.18E-27 | 3.04E-33 | 3.22E-21 | 2.21E-45 | 3.32E-08 |
| . | . | . | . | . | . | . | . | . | . |
| . | . | . | . | . | . | . | . | . | . |
| Proportion of stable sensor features per TW | 22.28% | 31.61% | 29.02% | 38.34% | 32.64% | 33.16% | 33.16% | 30.57% | 35.10% |
|  |  |  |  |  |  |  |  |  | Overall stability = 38.68% |

**Fig 10.**
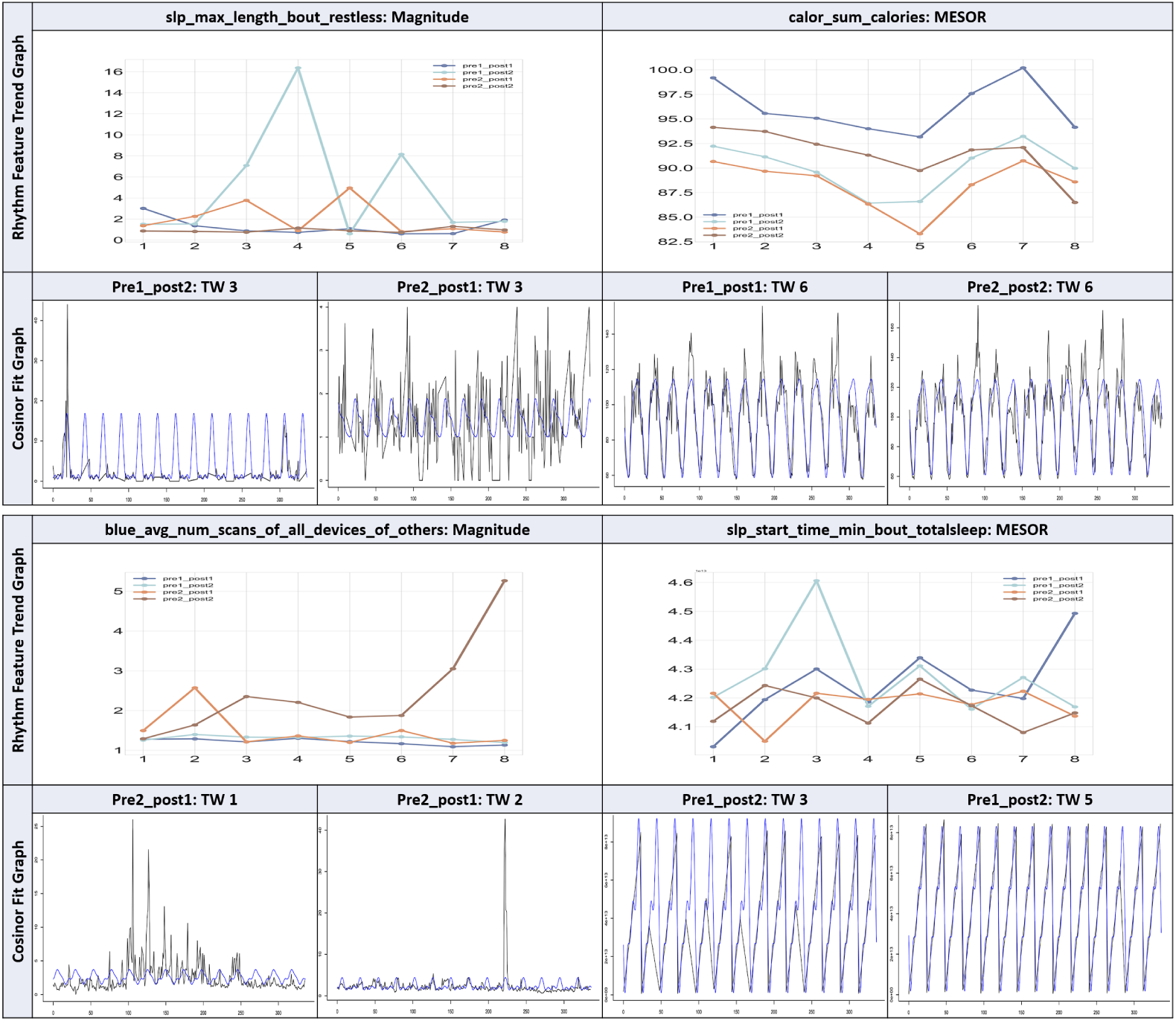
Loneliness cosinor rhythm features for four mental health groups during all time windows, and cosinor fit graph in single time window. This plot aggregates the cosinor results for four sensor features to show the stable patterns and two types of unstable patterns.

#### Prediction of Mental Health Status with Rhythmic Features

Our modeling of biobehavioral rhythms aims at observing cyclic behavior among students over the semester and to understand its potential relationship to students’ health status. From Autocorrelation, Periodogram, and Cosinor, we have seen that some groups have distinct characteristics. Following this thought, we can build classifiers to identify students’ mental health status according to their rhythms.

##### Dataset Generation

In our machine learning method, we use rhythmic features obtained from the Cosinor models as input to predict the post-semester loneliness and depression categories (low loneliness vs. high loneliness and no depression vs. with depression) of the students. We build two types of datasets, one with single sensors only and one with multiple sensors. For *Single Sensor* datasets, we use the rhythmic features of each sensor feature separately, *i*.*e*., for each sensor feature, and each time chunk takes the rhythmic features of this sensor feature and time chunk to form the input dataset.

We remove datasets with more than 30% missing instances (80 training instances) as we consider it too small to generate a reliable and generalizable model.

For *Multiple Sensors* datasets, we select the sensor features that give accuracy above baseline in models built with single sensors. We define baseline as the majority class, *i*.*e*., the category that has the highest percentage of labels for that category. We then repeat the same process we followed for single sensor datasets but this time for the combination of sensor features, *i*.*e*., for each time chunk and each combination of sensors, we take the rhythmic features of the selected sensor features of those sensors and time chunk to form the input dataset. Other than the input dataset’s difference, the machine learning pipeline is the same for the two types of datasets.

Given the imbalanced datasets for both health conditions *i*.*e*., different number of samples in the two classes (*e*.*g*., 59% of samples in category one vs. 41% in category 2 of depression), using the accuracy will not be adequate for performance evaluation and needs to be accompanied by other measures such as F1. For every combination of time window and sensor, the F1 score is used to select the model with the best performance. We build models with a single sensor and multiple sensors datasets for both mental health conditions. The results of all combinations are shown in Figures 11 and 12. The heatmaps use the depth of color to represent the F1 score. We only report results with accuracy above the baseline (majority class percentage). Through the single sensor modeling, we can judge which type of sensor is most effective in predicting mental health. By comparing the performance of modeling with a single sensor and modeling with multiple sensors, we find that the models with multiple sensors improve prediction performance. A summarization of the results are listed in Table 6.

**Table 6.** Summary of the mean and maximal values of F1 scorse for each combination of feature selection and machine learning methods shown in the heatmaps 11, 12. The bold values are either the biggest mean value of F1 scores, or the biggest maximal values of F1 scores

|  | Single Sensor |  |  |  |  |  | Multiple sensors |  |  |  |  |  |
| --- | --- | --- | --- | --- | --- | --- | --- | --- | --- | --- | --- | --- |
|  | Loneliness mean(max) |  |  | Depression mean(max) |  |  | Loneliness mean(max) |  |  | Depression mean(max) |  |  |
|  | GB | LR | RF | GB | LR | RF | GB | LR | RF | GB | LR | RF |
| IG | **0.69 (0.76)** | **0.69 (0.76)** | 0.66 (0.72) | **0.58 (0.70)** | **0.60 (0.61)** | 0.56 (0.63) | 0.73 (0.83) | 0.72 (0.78) | 0.69 (0.81) | 0.63 (0.83) | 0.60 (0.66) | 0.63 (0.76) |
| Lasso | 0.68 (0.72) | **0.70 (0.76)** | **0.74 (0.74)** | 0.57 (0.68) | 0.57 (0.64) | 0.55 (0.59) | 0.72 (0.78) | **0.75 (0.91)** |  | 0.59 (0.66) | **0.67 (0.89)** | 0.54 (0.54) |
| RLR |  | **0.70 (0.76)** | 0.68 (0.73) | 0.58 (0.65) | 0.56 (0.65) | 0.57 (0.60) | 0.75 (0.81) | 0.73 (0.82) | **0.76 (0.84)** | 0.65 (0.78) | 0.65 (0.79) | 0.65 (0.79) |

**Fig 11.**
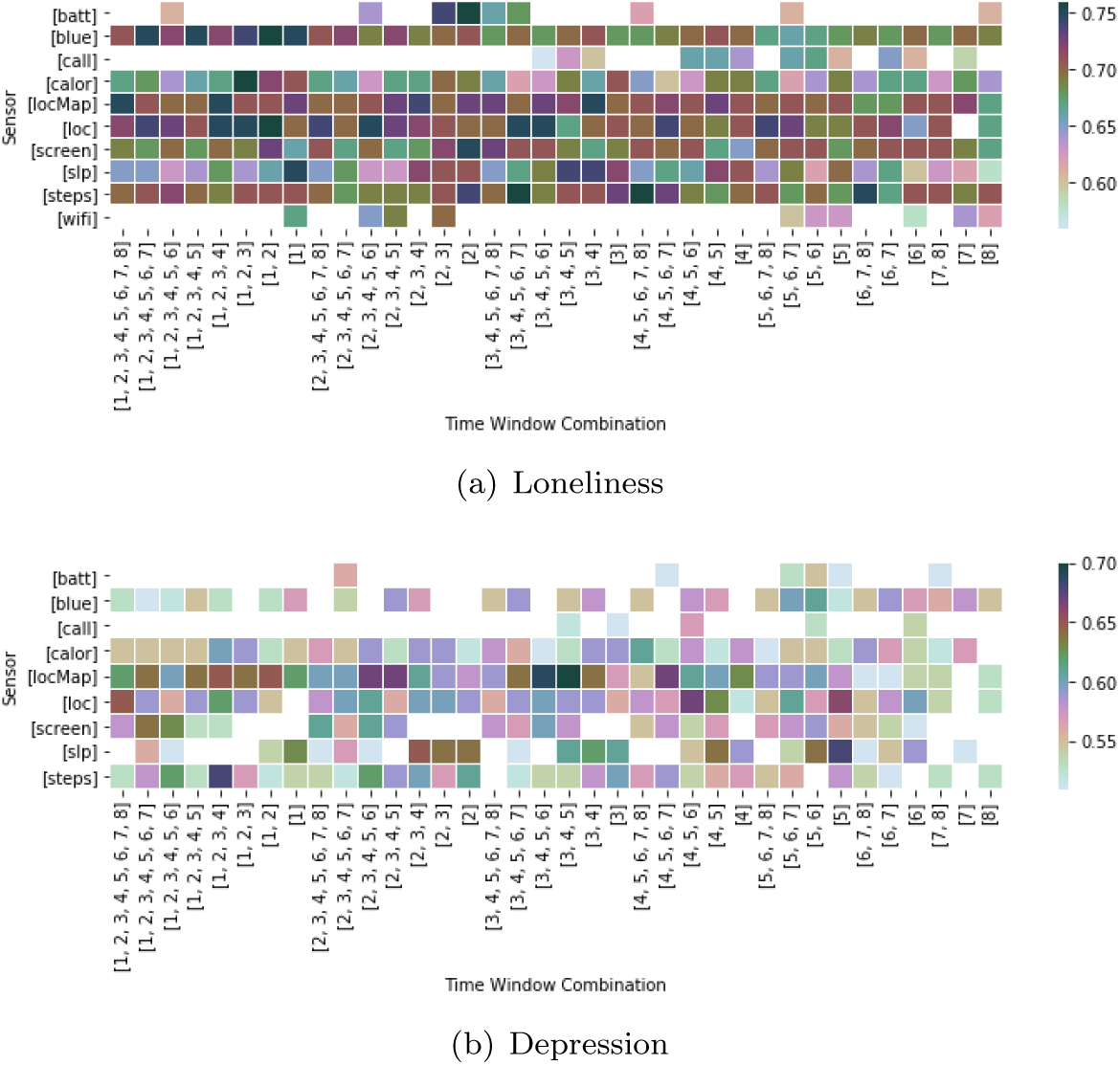
The heatmap displays the largest F1 score in the loneliness and depression prediction model trained by a combination of different single sensor features and time windows.

**Fig 12.**
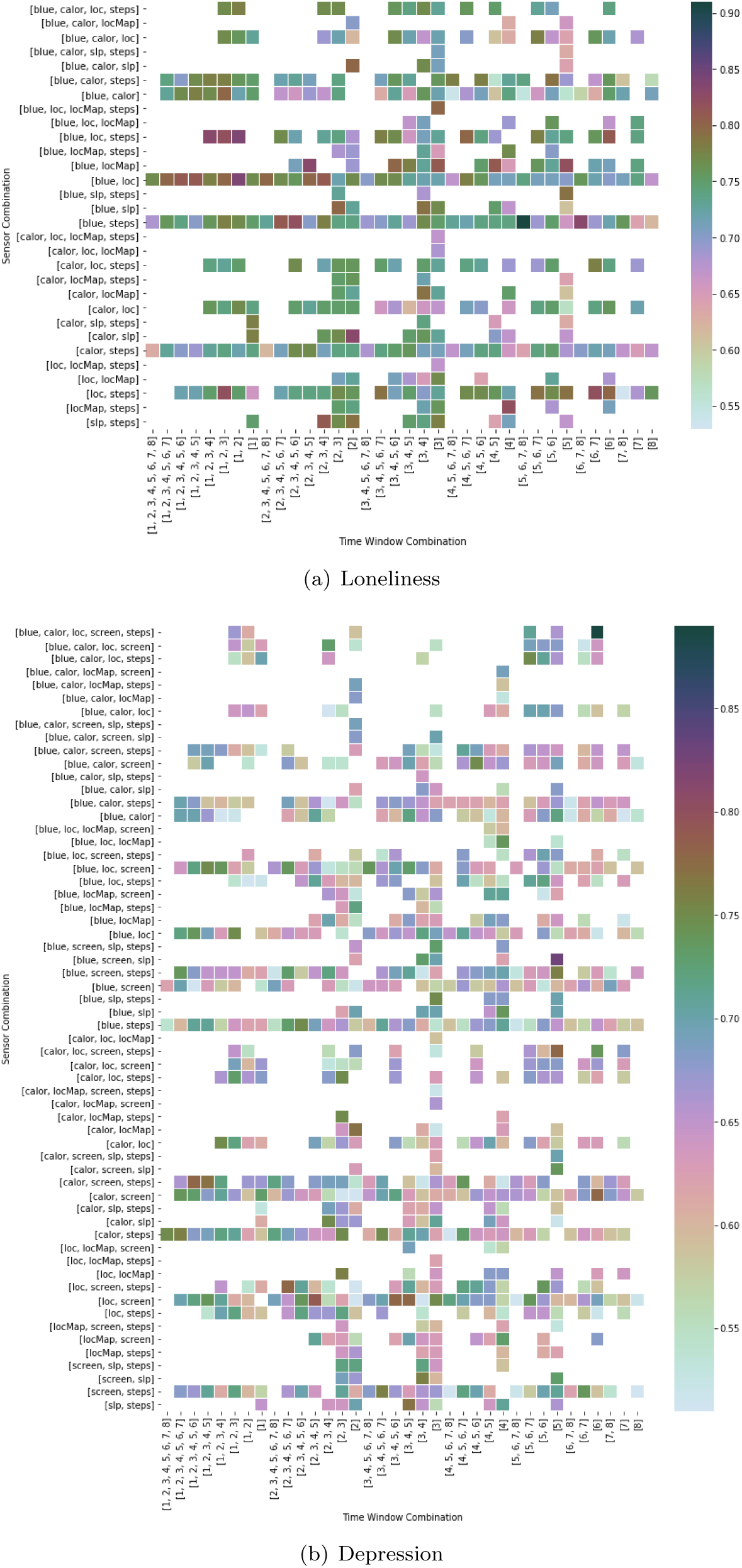
The heatmap displays the largest F1 score in the loneliness prediction model trained by a combination of different multiple sensor features and time windows.

##### Single Sensor Modeling

The F1 scores of machine learning models with single sensor features are shown in Figure 11. Rhythm parameters obtained from models built for features related to Bluetooth, calorie, location, sleep, and steps give better performance in predicting loneliness level. We observe that a similar set of features provides better results in depression prediction. Overall, the models for loneliness prediction obtain higher accuracy (F1) scores than depression models (Table 6). The best model to predict loneliness level is the Gradient Boosting model using the calorie data from *tw*_1_ to *tw*_3_ with an F1 score of 0.76. For depression, the highest F1 score is 0.7 obtained from the Gradient Boosting model using the locationMap data from *tw*_3_ to *tw*_5_. Compared to other sensors, models using rhythmic parameters from locationMap features show better performance for predicting depression (six out of ten models with the highest F1 score use locationMap features). Although the F1 scores of models with a single time window are generally lower than models with multiple time windows, there are some exceptions in the heatmaps of both loneliness and depression. For example, the loneliness model using the sleep features in *tw*_1_ achieves an F1 score of 0.75, and the F1 score of the depression model using the sleep features in *tw*_5_ equals 0.68.

##### Multiple Sensor Modeling

We do the same analysis for the combination of sensor features. From Figure 12, we observe that the combination of multiple sensor features contributes to the improvement of the F1 score. For example, the combinations related to steps, sleep, location, calorie, and Bluetooth end with better results. For predicting loneliness, the best model is built with Logistic Regression, which uses the Bluetooth and steps data from *tw*_5_ to *tw*_8_ and obtains an F1 score of 0.91. For predicting depression, the best model is obtained from Logistic Regression using the rhythm parameters from Bluetooth, calorie, location, screen, and step features. The model only uses *tw*_6_ to predict depression with an F1 score of 0.89. The best model predicting depression has a lower F1 score than the best model predicting loneliness, which is the same as the single sensor model.

Table 6 summarizes the mean and max of F1 scores for models built with each combination of feature selection and machine learning methods. In single sensor modeling, the combinations of Logistic Regression with Lasso and Randomized Logistic Regression) perform best for predicting loneliness with the mean and max F1 score of and 0.76, respectively. The combination of Gradient Boosting and Information Gain provides the highest F1 score for the prediction of depression. For the multiple sensor modeling, we observe that the maximum F1 scores of predicting loneliness and depression are 0.91 and 0.89, obtained from the combination of Logistic Regression and Lasso.

Overall our machine learning analysis demonstrates the ability of the framework to build massive micro-rhythmic models from each sensor feature and handle a broad set of features.

##### Dominant rhythm parameters that predict the mental health status

Although we used three feature selection methods in our evaluation, we observed that the Information Gain method provided more reliable and complete list of features during the training. Table 7 shows the rhythm features that are selected most frequently by Information Gain during depression prediction for each sensor feature in each time window. The vertical dominant feature (VDominant) is the most commonly selected feature for most of the sensors at a given time window, and the horizontal dominant feature (HDominant) is the most commonly selected feature in most time windows for a given sensor. The overall dominant feature (the feature at the bottom right corner in bold font) is the most commonly selected feature for all sensors and time windows. If two features are the most commonly selected features for the same number of sensors/time windows, we break the tie by taking the feature with higher frequency. Overall, Orthophase is selected most frequently for all sensors and time windows.

**Table 7.**
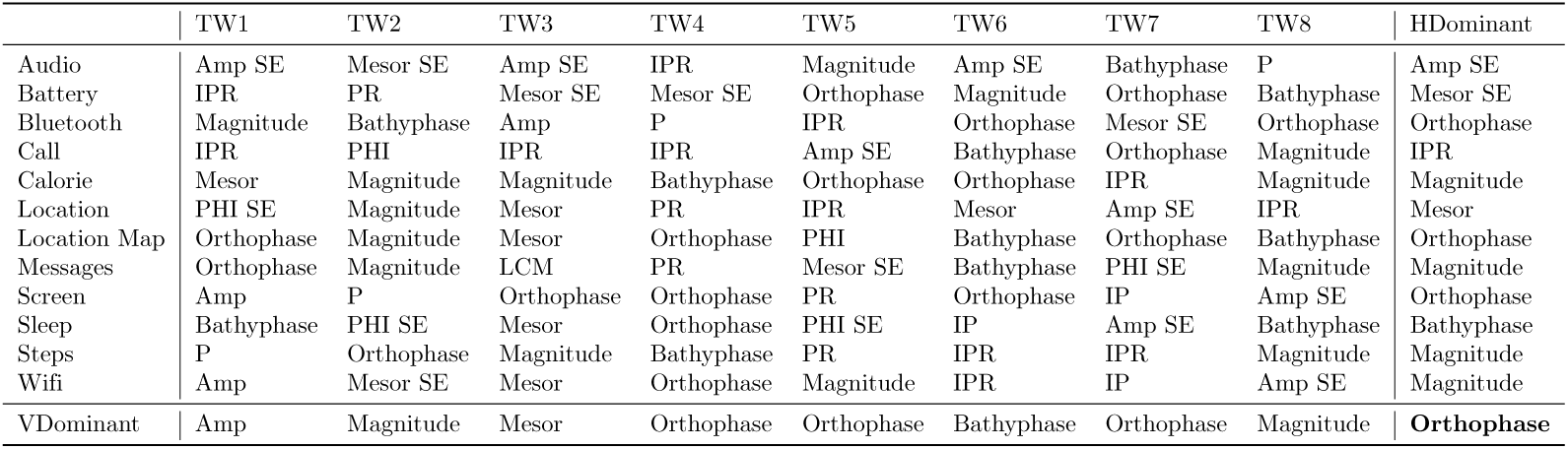
The most frequently selected rhythm features by Information Gain during depression prediction.

## Limitations and Future Work

Our computational framework for modeling biobehavioral rhythms enables the exploration of rhythmicity in time series data streams collected from mobile and wearable devices. It also provides the ability to explore further the impact of the rhythmic parameters obtained from cyclic models to predict an outcome of interest. Although, in this paper, only a set of specific functionalities and components are described as part of CoRhythMo, the framework is generalizable and can be adapted and extended to include more functionalities and features. The advancements include 1) adding more data sources such as weather, environment, work schedules, and social engagements to draw a more holistic picture of biobehavioral rhythms in individuals and groups of people, 2) adding a conclusive set of periodic functions and methods with diverse characteristics that provide the possibility of uncovering different cyclic aspects in data, and 3) advancing the machine learning component to incorporate a comprehensive selection of analytic methods that further enhance the capabilities of the framework to be used for predictive modeling of cyclic biobehavior.

For the current implementation, we limited our periodic functions to Autocorrelation, Periodogram, and Cosinor. In future work, we hope to build an ensemble system incorporating different types of rhythm detection algorithms and design a voting algorithm for aggregating the outputs of period detection algorithms. For example, the most frequently detected period by various detection algorithms will be treated as the dominant period.

A significant and novel contribution of this work is the proposal and application of the two rhythm stability measurement methods, namely CORESS and COSANOVA. As the central theme of this paper was introducing the computational framework, we did not do a detailed verification of the reliability of these two methods. However, we believe these methods can have a significant role in analyzing rhythmic behavior in time series data. So, we plan to rigorously evaluate these methods and introduce them to the research community as general methods for rhythm stability measurement.

So far, we have evaluated our framework with one dataset, and we realize that more experiments with various datasets are needed to thoroughly examine the capabilities of the framework and point out the necessary refinements. In this paper, we take mental health as an example to illustrate our computational framework’s feasibility in predicting health outcomes. However, we believe the framework can be used for many other outcomes, *e*.*g*., productivity, and lifestyle patterns. We are currently evaluating the framework in its ability to detect productivity cycles in college students and workers.

## Conclusion

We designed and presented a computational framework for modeling biobehavioral rhythms from mobile and wearable data streams that rigorously process sensor streams, detects periodicity in data, model rhythms from that data, and uses the cyclic model parameters to predict an outcome. Our evaluation of the framework using a dataset of smartphone and Fitbit data collected from college students over one semester showed that in addition to detection of rhythmicity, the framework could reliably discover various periods of different length in data, extract cyclic biobehavioral characteristics of students in different mental health groups through exhaustive modeling of rhythms for each sensor feature; and provide the ability to use a different combination of sensors and data features to predict their health status. The machine learning analysis for predicting mental health status demonstrated our framework’s ability to process a massive number of data streams to build and analyze micro-rhythmic models for each sensor feature and combinations of features and highlighted dominant rhythmic features for prediction of mental health status for each sensor across time windows. Our proposed methods for measuring rhythm stability in students also showed that different mental health groups project different rhythmic patterns over the course of the semester. These results provide valuable insights into studying students’ health status.

